# The Role of the Left Inferior Frontal Gyrus in Resolving Immediate and Carryover Semantic Interference During Picture Naming

**DOI:** 10.64898/2026.09.28.755022

**Authors:** Denise Y. Harvey, Harrison Stoll, Roy H. Hamilton

## Abstract

Naming pictures from the same semantic category can interfere with subsequent word retrieval. Although the left inferior frontal gyrus (LIFG) has been implicated in resolving this semantic interference, its causal role in this process remains unclear. Across three experiments, we investigated the contribution of the LIFG to interference resolution during blocked-cyclic naming. Experiment 1 tested whether continuous theta-burst stimulation over the LIFG increased semantic interference relative to vertex stimulation. Although the overall interference effect did not differ significantly between stimulation sites, exploratory analyses revealed an unexpected pattern based on the initial context (related vs. unrelated) with which items were named, suggesting that LIFG disruption affected retrieval of previously related items even when they were subsequently named in unrelated contexts. This observation motivated two further experiments examining the consequences of prior naming context more directly. Experiment 2 established that, without stimulation, interference observed in related blocks was no longer evident after the items were redistributed into unrelated blocks. Experiment 3 combined this design with cathodal or sham transcranial direct current stimulation (tDCS) over the LIFG. Planned comparisons showed that cathodal stimulation slowed naming of initially related items both within related blocks and after redistribution into unrelated blocks; corresponding comparisons for items named consistently in unrelated blocks were not significant. The similar patterns across two stimulation methods suggest that LIFG-mediated control may influence both selection under current competition and subsequent retrieval of previously competing words. These findings could be accommodated within a hybrid account in which working memory supports context-guided selection and selection-related control helps recalibrate lexical accessibility, limiting the carryover of interference into subsequent naming.

## Introduction

Fluent speech requires the rapid and accurate retrieval of intended words. Although this process typically occurs with little apparent effort, retrieving an intended word also activates unintended words that have similar meanings. Due to this coactivation, retrieving one word can interfere with the subsequent retrieval of semantically related words—a phenomenon termed semantic interference. The left inferior frontal gyrus (LIFG) has been implicated in supporting word retrieval in the presence of such interference, but its causal contribution remains incompletely understood. Determining causal role of the LIFG in resolving semantic interference is important for linking psycholinguistic accounts of word production to their neural implementation and may also illuminate why damage involving left frontal language regions—classically associated with Broca’s aphasia, a prototypical form of nonfluent aphasia—is accompanied by heightened susceptibility to semantic interference during naming (Biegler et al., 2008; Schnur et al., 2006; Arrigoni et al. 2024).

Psycholinguistic models of word production attribute semantic interference to processes involved in mapping semantic information onto lexical representations (Howard et al., 2006; Oppenheim et al., 2010; Roelofs, 2018). For example, when a speaker prepares to name a picture of a DOG, the pictured concept activates associated semantic features (e.g., has fur, has four legs), and this activation spreads to the corresponding lexical representation. Because some of these semantic features are shared with related concepts, such as CAT, lexical representations of semantically related, nontarget words also become activated. This coactivation can influence the efficiency of target-word retrieval, as evidenced by longer naming latencies when a picture is named after other semantically related pictures than when it is named after unrelated pictures (Belke, 2013; Damian & Als, 2005; Damian et al., 2001; Howard et al., 2006; Schnur, 2014). Importantly, the effects of prior naming can persist across successive naming trials. Two prominent computational models explain these lasting effects in terms of incremental changes to the connections between semantic and lexical representations (Howard et al., 2006; Oppenheim et al., 2010). Although the models differ in the nature of the proposed changes and in whether lexical selection is assumed to be competitive, both predict that naming produces lasting changes within the lexical-semantic system that affect the subsequent retrieval of semantically related words.

The blocked-cyclic naming task provides a means of examining these lasting effects across repeated retrievals (e.g., Damian et al., 2001). In this task, participants repeatedly name small sets of pictures presented in blocks containing either semantically related or unrelated pictures. Each block consists of multiple presentation cycles, with every picture in the set appearing once per cycle. Naming latencies typically decrease across cycles in unrelated blocks, reflecting facilitation from repeated retrieval (i.e., repetition priming). In related blocks, however, this facilitation is reduced or offset by semantic interference, resulting in longer naming latencies in the related than the unrelated condition after the first presentation cycle (hereafter, *semantic blocking effect*; e.g., Belke et al., 2005; Belke, 2017). Because the same pictures are named in both related and unrelated blocks, and interference emerges only after the first presentation cycle, the task provides a controlled means of examining the neural processes associated with semantic interference while holding the target pictures and general naming demands constant.

The LIFG has been implicated in selecting among coactivated representations and supporting word retrieval under conditions of semantic interference (Arrigoni et al. 2024; Pino et al., 2022; Riès et al., 2017). Across language production tasks, greater LIFG recruitment is observed when selection demands are high, including when speakers must generate a verb in response to nouns associated with multiple plausible responses (e.g., Snyder et al., 2011). Evidence specific to blocked-cyclic naming comes from converging neuroimaging and neuropsychological findings. In neurologically healthy adults, LIFG activation was greater during naming in semantically related than unrelated blocks. Moreover, the magnitude of this activation difference positively correlated with the number of errors produced in related blocks, such that individuals who experienced greater semantic interference also showed greater LIFG recruitment (Schnur et al., 2009). Complementary neuropsychological evidence demonstrated that, compared with persons with fluent aphasia and healthy older controls, persons with nonfluent aphasia exhibited slower and more error-prone naming in semantically related than unrelated blocks (Schnur et al., 2006). In a subsequent lesion analysis, greater interference-related error rates were associated with damage to the LIFG (Schnur et al., 2009). However, neuroimaging evidence is correlational, and stroke lesions are heterogeneous, often extending beyond a single cortical region. Indeed, lesion and diffusion-imaging evidence indicates that susceptibility to semantic interference is also related to temporal-lobe damage and the integrity of underlying white-matter pathways (Harvey & Schnur, 2015). Thus, although these findings implicate the LIFG in word retrieval under semantic interference, they do not fully isolate its contribution or establish the precise process reflected by its recruitment.

Noninvasive brain stimulation offers a complementary approach by transiently altering LIFG function within an otherwise intact language system, thereby allowing its contribution to semantic interference to be tested more directly. To date, however, causal evidence specific to semantic interference in cyclical naming remains limited. Krieger-Redwood and Jefferies (2014) found that offline, low-frequency (1 Hz) repetitive transcranial magnetic stimulation (TMS) intended to disrupt LIFG function reduced semantic facilitation during the first presentation cycle but did not alter semantic interference during later cycles. Thus, although their findings demonstrated a causal contribution of the LIFG to semantically driven word retrieval, whether the LIFG is necessary for resolving the semantic interference that emerges with repeated naming remains unclear.

The present study examined this question across three experiments. In Experiment 1, we applied continuous theta-burst stimulation (cTBS)—a noninvasive brain stimulation (NIBS) protocol commonly used to transiently reduce cortical excitability—to the LIFG of neurologically healthy adults to perturb its function during blocked-cyclic naming. This approach allowed us to examine the consequences of transiently disrupting LIFG function within an otherwise intact language system. Prior work using anodal transcranial direct current stimulation (tDCS), a protocol commonly associated with increased cortical excitability, over the LIFG reduced the semantic blocking effect, suggesting that modulating LIFG function via NIBS can alter semantic interference during naming (Pisoni et al., 2012). Participants completed the blocked-cyclic naming task in a within-subject design, once following cTBS to the LIFG and once following cTBS to the vertex as a control site. If the LIFG is necessary for resolving semantic interference during word retrieval, then perturbing its function should produce a larger semantic blocking effect following LIFG than vertex stimulation. Exploratory analyses of Experiment 1 revealed that the effects of LIFG stimulation differed depending on the context (related vs. unrelated) with which items were first presented for naming. This unexpected pattern motivated two follow-up experiments designed to clarify its source: a behavioral study without brain stimulation (Experiment 2) and a study using cathodal tDCS over the LIFG (Experiment 3).

## Experiment 1: cTBS Study

### Overview

Experiment 1 used a within-subject, active-control design to test the causal contribution of the LIFG to semantic interference during blocked-cyclic naming. Neurologically healthy adults completed two experimental sessions in which continuous theta-burst stimulation (cTBS) was applied to either the LIFG or the vertex as a control site. Before cTBS, participants completed a verb-generation task adapted from Snyder et al. (2011), which engages LIFG-supported processes involved in retrieving and selecting words among alternative responses. Because the effects of NIBS depend partly on the activation state of the targeted neural populations, the verb-generation task was administered to engage the relevant language network and standardize its cognitive state before stimulation (Silvanto & Pascual-Leone, 2008). Following cTBS, participants completed the blocked-cyclic naming task.

### Participants

Twenty participants (9 female; mean (standard deviation [SD]) age = 26 [7.04]) recruited from the University of Pennsylvania community participated in Experiment 1. All participants were native English speakers, had normal or corrected-to-normal vision, and provided informed consent in accordance with the University of Pennsylvania Institutional Review Board.

### Materials & Design

*Verb generation task.* Stimuli were 100 written nouns with either high or low association strengths with possible verb responses (retrieval demand) crossed with either high or low competition with alternative verb responses (25 per condition). To ensure that different nouns were presented in each of the two stimulation sessions, stimuli were divided into two lists of 50 nouns each, which were matched on association strength (*t*(98) = .89, *p* .87) and competition strength (*t*(98) = 1.59, *p* = .11).^1^ Mix software (van Casteren & Davis, 2006) was used to generate different trial orders per list for each participant with the constraint that no more than three consecutive trials presented a noun from the same condition.

#### Blocked-cyclic naming task

Stimuli were 144 colored photographs depicting familiar objects taken from various image databases and Internet sources (Adlington et al., 2009; Brodeur et al., 2010; Guérard, & Bouras, 2014; Kovalenko et al., 2012; Moreno-Martínez & Montoro, 2012; Viggiano et al., 2004). There were 12 semantic categories consisting of 12 exemplars each. To ensure that different items were presented each of the two stimulation sessions, category exemplars were split into two lists containing six items per category for a total of 72 items per list (see Appendix A). The lists were matched for name agreement, lexical frequency, visual complexity, and number of syllables (*p*’s > .22).

Within each list, there were 12 related blocks comprising of 6 (or half) of the exemplars in each category. To create the unrelated blocks, one item from each of six categories formed six 6-item unrelated sets, and one item from each of the remaining six categories formed an additional six 6-item unrelated sets, forming a total of 12 unrelated blocks. Each list contained 24 blocks. Within a block, the 6-item sets were presented in pseudorandom order for four cycles (24 trials), for a total of 576 trials within a single experimental session. Thus, each item appeared an equal number of times in the related and unrelated sets.

Four orders of each 72-item list were generated using Mix software such that no more than three related or unrelated blocks appeared consecutively and consecutive trials within and across blocks did not consist of the same object or phonological onset (van Casteren & Davis, 2006). The first 12 blocks (hereafter, *first half*) consisted of six related and six unrelated sets that did not contain any of the same items. In the remaining 12 blocks (hereafter, *second half*), items previously presented in the unrelated sets formed the related sets and vice versa (see Figure 1). We counterbalanced the related and unrelated sets appearing in each half across participants.

**Figure 1.**
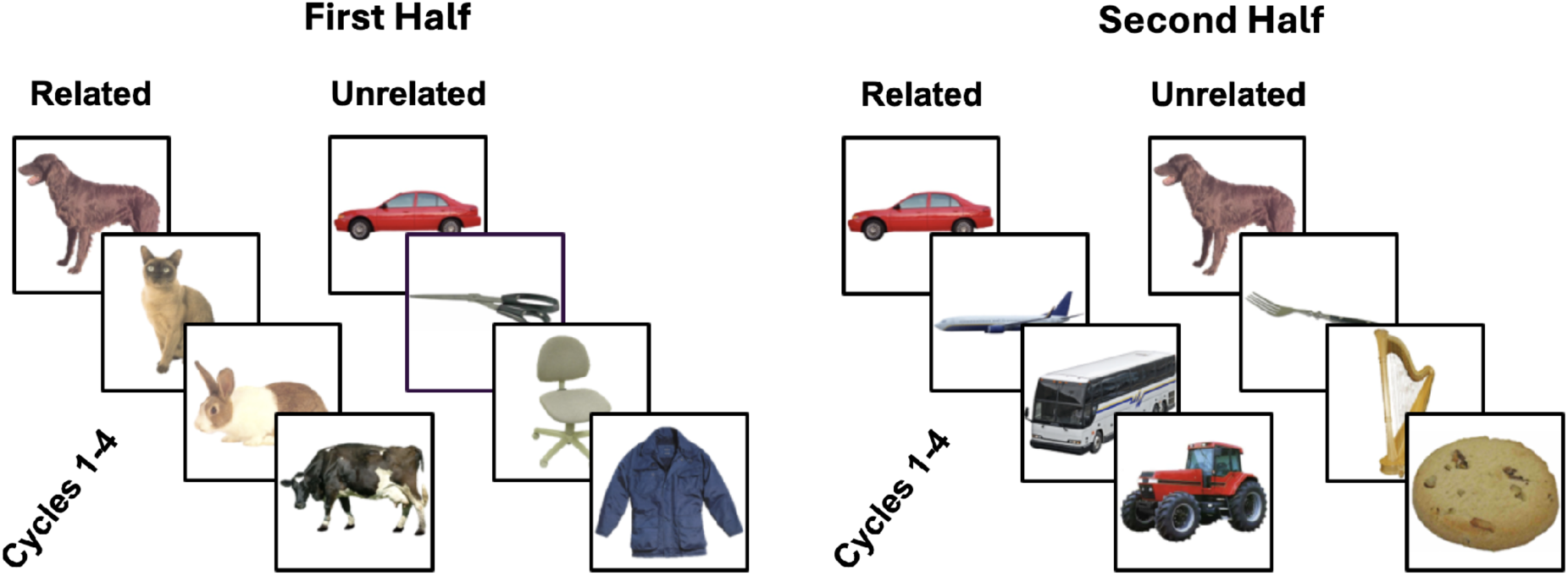
Illustration of the blocked-cyclic naming task for Experiment 1. Pictures appeared in sets of semantically related and unrelated contexts. In the first half, four items were repeated across four cycles in either related or unrelated sets (left panel). In the second half, these same items were presented in the opposite context (right panel): items named previously in unrelated sets were reorganized into related sets and vice versa.

### Apparatus

The experiment was presented on a laptop computer using E-Prime software. A microphone connected to E-Prime SRBox detected vocal responses using a voice key trigger to record response times (RTs) to the nearest millisecond (ms). A separate digital recorder captured verbal responses for later review of response accuracy and to recalculate RTs when the voice key was mistriggered by an extraneous sound.

### Procedure

Participants attended two sessions: one in which they received cTBS stimulation to the LIFG, and another where stimulation was delivered to the vertex. The two sessions were separated by at least 7 days. For each participant, we presented different stimulus lists (List 1 or List 2 for the blocked-cyclic naming and verb generation tasks) in each session. Each session lasted about 45 minutes and followed this order: blocked-cyclic naming stimuli familiarization, verb generation task, cTBS, and blocked-cyclic naming task.

To familiarize participants with the blocked-cyclic naming stimuli, each of the 72 objects was presented centrally on the screen with its typed name displayed underneath the picture. The picture and its written label (or name) remained on the screen until the participant pressed the spacebar, indicating they understood the correct response for that object. Participants were instructed to refrain from saying the image’s name out loud. This process continued until participants had viewed and acknowledged all 72 objects.

After the familiarization phase and prior to stimulation, participants performed the verb generation task. This was done to engage the language/control network targeted for stimulation prior to receiving stimulation, which is thought to enhance and/or localize the effects of stimulation (e.g., Gervits et al., 2016; Gill et al., 2015). Participants were presented with text specifying a noun and instructed to produce the first verb that came to mind; the verb must either be something the noun does or something that can be done with the noun (following Snyder & Munakata, 2008). Participants were then given an example (e.g., in response to the noun GRASS, one could say “grow” or “mow”) followed by five practice trials of noun cues not used in the actual experiment. Nouns remained on the screen for 5,000 ms or until the voice key was triggered. In between each trial, a fixation cross “+” appeared in the center of the screen for 1,000 ms. The task lasted approximately 5-7 minutes.

Immediately following stimulation, participants performed the blocked-cyclic naming task. Before the start of the task, participants were reminded to use the names provided in the familiarization phase for their response. Each picture was presented centrally on the screen, and participants were instructed to name the picture as quickly and accurately as possible. The onset of picture presentation coincided with a “beep” to indicate the trial onset. Each session was also recorded on a separate recorder, which was later used to check response accuracy and manually recalculate the RTs for responses that were not triggered by the voice key. Like the verb generation task, there was a 5,000 ms response deadline and 1,000 ms inter-stimulus interval (ISI) of 1,000 ms displaying a fixation (+). The task lasted approximately 25 minutes.

### Stimulation parameters

Stimulation was administered using a 70 mm diameter hand-held figure-of-eight coil (Magstim Super Rapid 2 Plus 1 Transcranial Magnetic Stimulator; Magstim, Whitland, UK). Prior to stimulation, we obtained high-resolution structural MRI scans for each participant. The Brainsight (Rogue Inc., Montreal, Canada) neuronavigational system was used to mark targets for stimulation (in native space), and co-register structural MRI volumes with respect to the location of the participant and the coil. Participants received cTBS to the LIFG and vertex in separate sessions. The cTBS protocol entailed continuous delivery of 50 Hz triplets of TMS pulses at 5 Hz for a total of 600 pulses (40 s). Following prior work, we delivered stimulation at 80% of active motor threshold (aMT), as defined using standard methods (see e.g., Huang et al., 2005). AMT was established in a separate session as part of a larger study (REFs). The experimental cTBS target was the anterior portion of the LIFG, i.e., the left pars triangularis (lPTr; e.g., Naeser et al., 2005a, Barwood et al., 2012). The vertex (control) cTBS target was located at the midline between the primary somatosensory and motor cortices. We randomized the stimulation-site session order, with 10 participants receiving LIFG stimulation first and 10 receiving vertex stimulation first.

### Data Analysis

Trial-level naming latencies were analyzed using linear mixed-effects regression in R with the *lme4* package (Bates et al., 2015). The initial model included fixed effects of Stimulation Site (LIFG vs. vertex), Condition (related vs. unrelated), and their interaction. The random-effects structure included by-participant random intercepts and slopes for Session, with the correlation between the intercept and slope estimated, and item random intercepts. The model was fit using the *bobyqa* optimizer with a maximum of 200000 iterations.

We excluded trials containing incorrect responses, voice-key errors, or naming latencies shorter than 250 ms consistent with prior semantic-interference and picture-naming studies (Riley et al., 2015; see also Belke, 2013; Schnur et al., 2006 for a similar approach).^2^ We also excluded cycle 1 because semantic interference typically emerges after the initial presentation cycle and remains relatively stable across subsequent cycles (Belke & Stielow, 2013; Belke et al., 2005; Belke, 2017). Fixed effects were evaluated using Type III F tests with Satterthwaite-approximated degrees of freedom (*lmerTest* package; Kuznetsova et al., 2017). Simple-effects contrasts of estimated marginal means were evaluated using asymptotic z tests (*emmeans* package; Lenth, 2023). Holm-adjusted p-values were calculated across the two condition contrasts (one per stimulation site) and, separately, across the two stimulation-site contrasts (one per condition). We evaluated statistical significance at α = .05.

### Results

A total of 3.72% of data points in cycles 2-4 (642) were excluded from the analysis, which comprised of incorrect responses or false starts (3.40%) and equipment or microphone errors (0.31%).

Full model results are reported in Table 1. The results revealed a significant main effect of condition (*F*(1, 16454.85) = 127.5, *p* <. 001) and no significant main effect of Stimulation Site (*F*(1, 19.27) = 1.71, *p* = .20); however, the Stimulation Site x Condition interaction did not reach significance, *F*(1, 16454.86) = 3.44, *p* = .06. The simple effects contrasts found that participants were significantly faster to name pictures presented in unrelated vs. related contexts following cTBS to the LIFG *(b* = -24.6, *z* = -6.67, *p* < .001) and vertex (*b* = -34.2, *z* = -9.31, *p* < .001). However, naming latencies did not differ as a function of Stimulation Site for items presented in related contexts (*b* = -13.3, *z* = -.95, *p* = .35) or unrelated contexts (*b* = -23.0, *z* = -1.63, *p* = .21).

**Table 1.** Experiment 1: Type III F Tests.

Experiment 1: Type III F Tests
| Effect | SS | MS | $df_1$ | $df_2$ | F | p |
| --- | --- | --- | --- | --- | --- | --- |
| Stimulation Site | 48288.00 | 48288.00 | 1 | 19.30 | 1.71 | .206 |
| Condition | 3593636.00 | 3593636.00 | 1 | 16454.80 | 127.54 | <.001 |
| Stimulation Site ×<br>Condition | 97020.00 | 97020.00 | 1 | 16454.90 | 3.44 | .064 |

*Note.* SS = sum of squares; MS = mean square; $df_1$ = numerator degrees of freedom; $df_2$ = denominator degrees of freedom, computed using Satterthwaite's approximation. × denotes an interaction term.

**Table 2.**
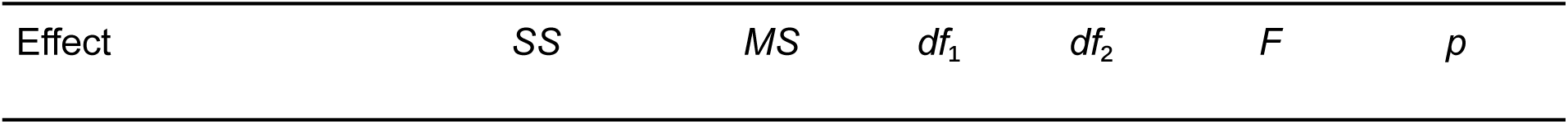
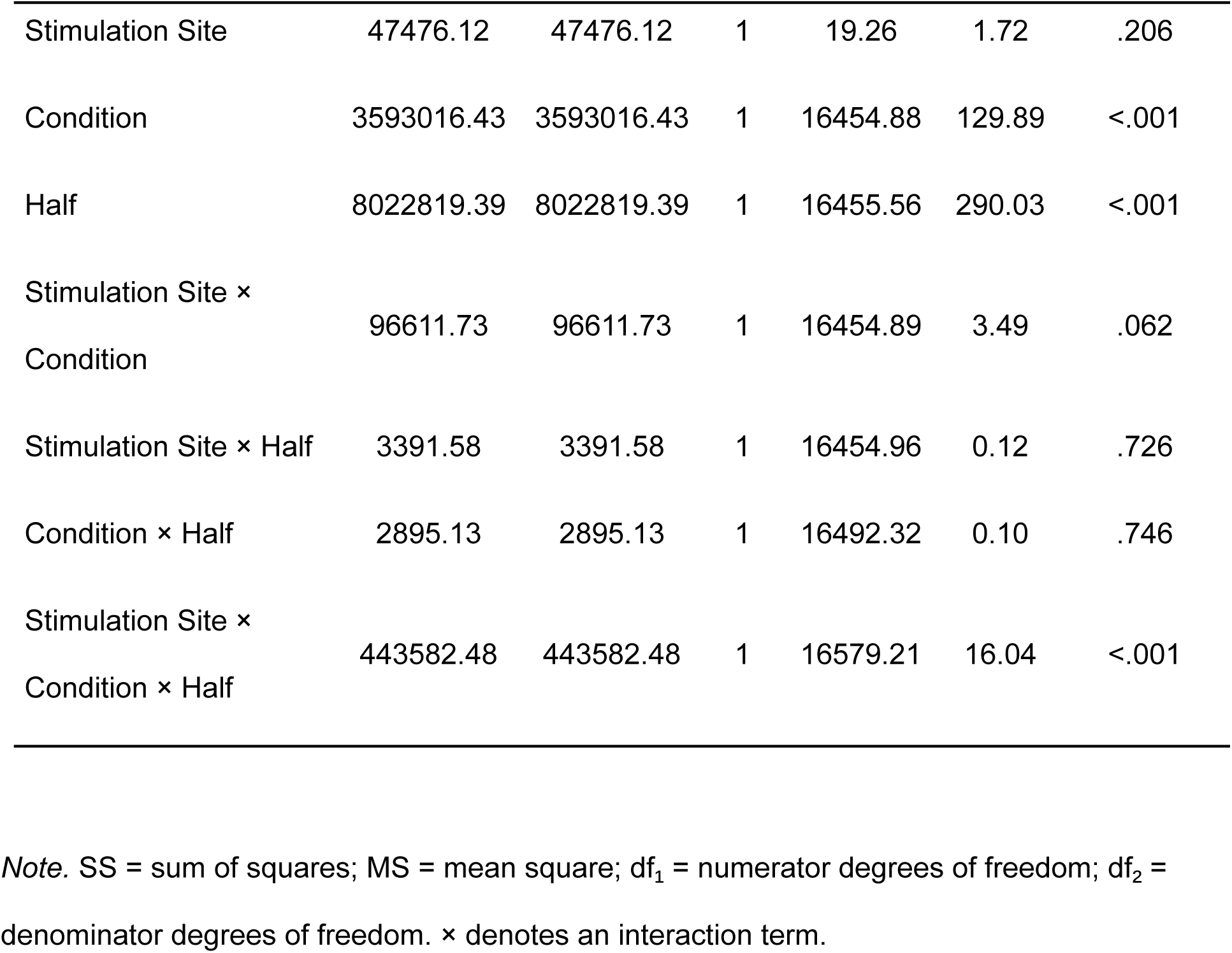
Experiment 1: Type III F Tests (Includes Three-way Interaction Term with Half)

| Effect | SS | MS | df<sub>1</sub> | df<sub>2</sub> | F | p |
| --- | --- | --- | --- | --- | --- | --- |
| Stimulation Site | 47476.12 | 47476.12 | 1 | 19.26 | 1.72 | .206 |
| Condition | 3593016.43 | 3593016.43 | 1 | 16454.88 | 129.89 | <.001 |
| Half | 8022819.39 | 8022819.39 | 1 | 16455.56 | 290.03 | <.001 |
| Stimulation Site ×<br>Condition | 96611.73 | 96611.73 | 1 | 16454.89 | 3.49 | .062 |
| Stimulation Site × Half | 3391.58 | 3391.58 | 1 | 16454.96 | 0.12 | .726 |
| Condition × Half | 2895.13 | 2895.13 | 1 | 16492.32 | 0.10 | .746 |
| Stimulation Site ×<br>Condition × Half | 443582.48 | 443582.48 | 1 | 16579.21 | 16.04 | <.001 |

*Note.* SS = sum of squares; MS = mean square; $df_1$ = numerator degrees of freedom; $df_2$ = denominator degrees of freedom. × denotes an interaction term.

Contrary to our hypothesis, LIFG stimulation did not significantly increase the semantic blocking effect relative to vertex stimulation. However, the primary analysis collapsed across task position and therefore could not capture potential changes in the stimulation effect over the experiment. Because the effects of prior naming are known to persist across subsequent trials and blocks, we conducted an exploratory analysis to determine whether the effects of Stimulation Site and Condition depended on the order in which items were encountered in related versus unrelated contexts over the course of the task. We expanded the original mixed-effects model to include Half (first vs. second) and its interactions with Stimulation Site and Condition. The critical test was the Stimulation Site × Condition × Half interaction.

As shown in Figure 2, we found a statistically significant three-way interaction between Stimulation Site, Condition, and Half (*F*(1, 16579.21) = 16,04, *p* < .001). Follow-up contrasts showed the expected blocking effect in the first half of the task, with slower naming of related items after LIFG stimulation relative to Unrelated items (*b* = -34.2, *z* = -6.59, *p* < .001). This effect was also found in the vertex stimulation (*b* = -22.9, *z* = -4.43, *p* < .001). Critically, we found that the LIFG was slower than the vertex condition for naming in the related condition *(b=* -24.71, *z* = -1.69, *p* = .09). Surprisingly, for the second half, we found that naming latencies were slower for unrelated items following LIFG vs. vertex stimulation (*b =* -32.54, *z* = -2.32, *p* < .05). However, naming latencies for the related condition did not differ across sites *(b =* -2.01, *z* = -.14, *p* = .89).

**Figure 2.**
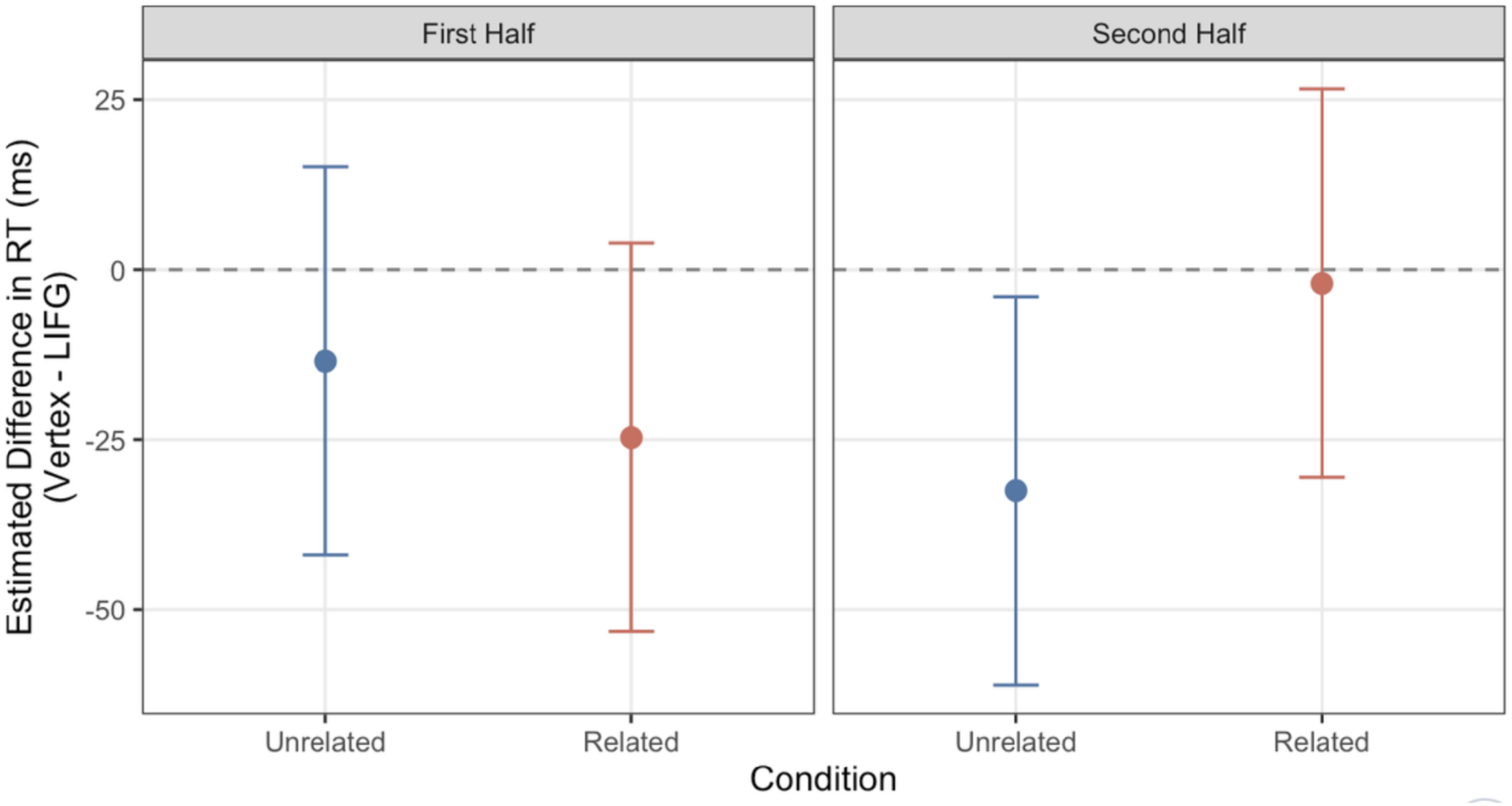
Model-estimated RT for Experiment 1. Values below the dotted line indicate faster responses during vertex stimulation than LIFG stimulation.

### Experiment 1 Summary

Experiment 1 provided preliminary evidence that perturbing LIFG function had different consequences across the two halves of the task. In the first half, we found a trend toward a difference between the LIFG and the vertex for related pictures. Surprisingly, in the second half, this effect emerged for pictures presented in unrelated contexts, suggesting that interference experienced earlier in the task may have had downstream consequences even after the semantic context changed. Because this pattern emerged from exploratory analyses, Experiment 2 tested whether the corresponding change across task halves emerges behaviorally in the absence of brain stimulation.

## Experiment 2: Behavioral Study

### Methods

#### Overview

Experiment 2 employed a behavioral-only design and a modified blocked-cyclic naming task to investigate whether the pattern observed in Experiment 1 reflected the lasting influence of the semantic context in which items were initially named or if it was the result of inhibiting the LIFG. In this version of the task, all items in the second half of the study belonged to unrelated sets. Items initially named in unrelated contexts remained in unrelated contexts (’unrelated-unrelated’ [UU]), whereas items initially named in related contexts were redistributed into blocks consisting of unrelated contexts (’related-unrelated’ [RU]; see Figure 3). This design allowed us to test whether prior naming in a related context continued to affect performance when the items were subsequently named in an unrelated context. Unlike Experiment 1, participants did not perform the verb generation task, and we did not administer brain stimulation.

**Figure 3.**
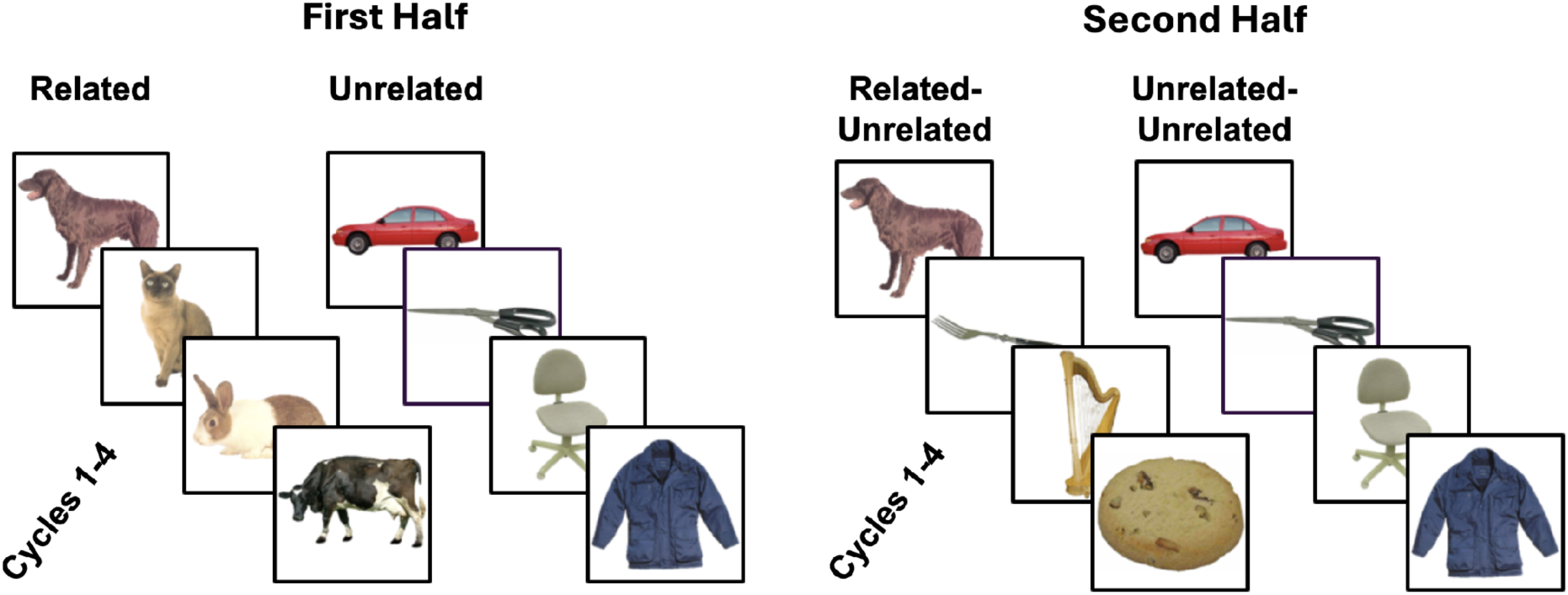
Illustration of the blocked-cyclic naming task for Experiments 2 and 3. Pictures appeared in sets of semantically related and unrelated contexts. In the first half, four items were repeated across four cycles in the related and unrelated sets (left panel). In the second half, the related items were rearranged to unrelated sets (related-unrelated sets), and the unrelated items were kept in the same sets (unrelated-unrelated sets).

#### Participants

Twenty-one healthy college students were recruited from the University of Pennsylvania to participate in Experiment 2. None of the participants were bilingual, and all participants were native English speakers with normal or corrected to normal vision. All participants gave informed consent in accordance with the protocol approved by the University of Pennsylvania IRB. Data from one participant was dropped from analyses due to high error rate (> 20% trials). Data submitted for analyses came from 20 participants (13 females, mean [SD] age = 20 [1.17]).

#### Design

##### Blocked-cyclic naming

The task design followed that of Experiment 1 with the following exception: In the second half of the experiment, all items were presented in unrelated contexts. This meant that one item from each of the 6 six-item sets presented in related blocks in the first half of the experiment comprised 6 six-item additional unrelated blocks presented in the second half of experiment (see Figure 3). As in Experiment 1, participants had a response deadline of 5,000 ms with an ISI of 1,000 ms.

#### Procedure

As in Experiment 1, participants were first familiarized with the pictures and their intended names using the same procedure. They then completed the blocked-cyclic naming task according to the procedures described for Experiment 1. The verb-generation task was not administered because Experiment 2 did not involve brain stimulation and therefore did not require prior engagement of a targeted language network.

#### Analysis

Trial-level naming latencies were analyzed using a linear mixed-effects model following the procedures described for Experiment 1. The model included Context Sequence (related-to-unrelated [RU] vs. unrelated-to-unrelated [UU]), Half (first vs. second), and their interaction as fixed effects. The Context Sequence × Half interaction tested whether the effect of initial semantic context differed between the two halves of the task. The model included by-participant random intercepts and item random intercepts. Trial-exclusion criteria and all other analytic procedures were identical to those used in Experiment 1, with the additional exclusion of RTs >2,000 ms because, in the absence of stimulation, these unusually long responses were considered more likely to reflect outliers.

#### Results

Of the cycle 2-4 trials, a total of 3.45% data points (298) were excluded from analysis due to comprising of incorrect responses or false starts (2.87%), equipment or microphone errors (0.47%), or outliers (0.10%).

We found a statistically significant Half x Context Sequence interaction (*F*(1, 8251.77) = 60.71, *p* < .001; see Figure 4). Simple-effects contrast tests examined differences across conditions within each half. In the first half, before the semantic contexts changed, participants named items presented in unrelated contexts faster than those named in related contexts, demonstrating the expected semantic blocking effect (*b* = -47.5, *z* = -9.59, *p* < .001). In the second half—when all pictures were presented in unrelated contexts with the critical difference being the prior context with which they were presented for naming—naming latencies did not differ between the RU and UU conditions, *b* = 6.94, *z* = 1.41, *p* = .16, indicating no detectable effect of prior related-context retrieval. See Table 3 for full model results.

**Figure 4.**
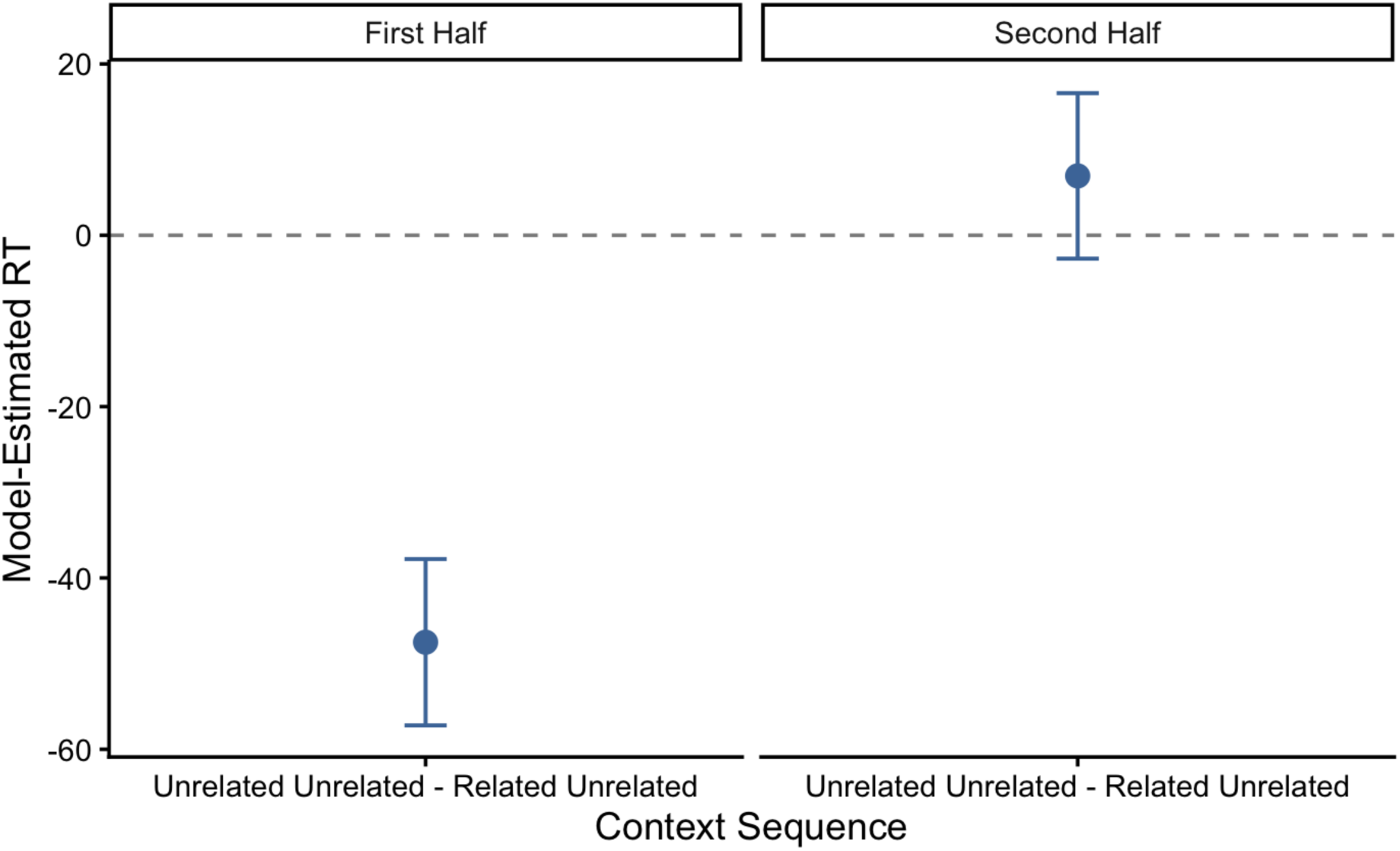
Model-estimated RT for Experiment 2. The dotted line represents no difference between each context condition. Thus, values below the dotted line represent faster naming latencies for items presented in unrelated contexts (i.e., the semantic blocking effect), and positive values represent no difference in naming latencies based on the semantic context).

**Table 3.**
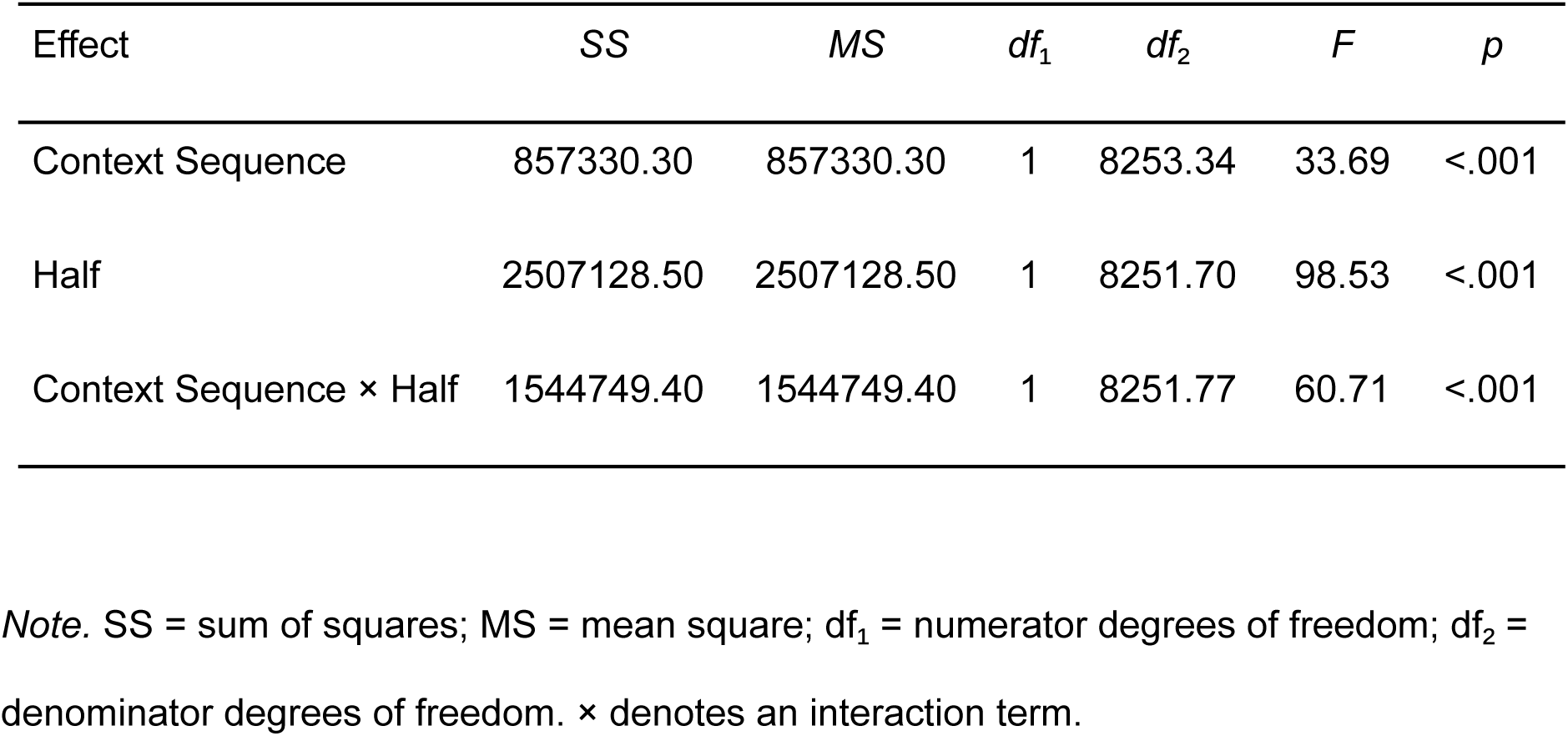
Experiment 2: Type III F Tests.

| Effect | SS | MS | $df_1$ | $df_2$ | $F$ | $p$ |
| --- | --- | --- | --- | --- | --- | --- |
| Context Sequence | 857330.30 | 857330.30 | 1 | 8253.34 | 33.69 | <.001 |
| Half | 2507128.50 | 2507128.50 | 1 | 8251.70 | 98.53 | <.001 |
| Context Sequence × Half | 1544749.40 | 1544749.40 | 1 | 8251.77 | 60.71 | <.001 |

*Note.* SS = sum of squares; MS = mean square; $df_1$ = numerator degrees of freedom; $df_2$ = denominator degrees of freedom. × denotes an interaction term.

#### Experiment 2 Summary

The first-half semantic blocking effect confirmed that the paradigm induced semantic interference in this sample. In the second half, however, naming latencies did not differ as a function of the semantic context in which pictures were initially presented. Thus, in the absence of perturbation to the LIFG, we found no evidence of semantic interference carryover. This pattern of results contrasts with the exploratory findings from Experiment 1, in which perturbation of the LIFG affected naming pictures in unrelated contexts when they were previously named in related contexts. Experiment 3 therefore tested whether modulating LIFG function with tDCS, a physiologically distinct form of NIBS commonly associated with reduced cortical excitability, would produce a similar pattern. Converging findings across stimulation methods alongside the modified blocked-cyclic naming design would strengthen the interpretation that the LIFG contributes to limiting the effects of prior semantic interference.

## Experiment 3: Cathodal tDCS Study

### Methods

#### Overview

Similar to Experiment 1, Experiment 3 used a within-subject, active-sham design to test the causal contribution of the LIFG to semantic interference. However, here, we used the modified blocked-cyclic naming task implemented in Experiment 2. Neurologically healthy adults completed two experimental sessions in which cathodal or sham tDCS was applied to the LIFG. As in Experiment 1, participants completed a verb generation task prior to stimulation in order to engage the language network and therefore increase the likelihood of tDCS-induced neuromodulatory effects on word retrieval performance.

#### Participants

Twenty-one healthy participants were recruited through the University of Pennsylvania to participate in Experiment 3. Inclusion criteria followed that of the previous experiments. Thus, one participant was excluded due to being left-handed, another due to being a non-native English speaker, and two were excluded due to equipment errors. All participants gave informed consent in accordance with the protocol approved by the University of Pennsylvania IRB. Data submitted from analyses came from 16 participants (11 females, mean [SD] age = 21.87 [4.77]).

#### Design

##### Verb generation

The verb generation task administered in Experiment 3 was identical to that of Experiment 1, again with the goal of engaging the language network, potentially increasing NIBS effectiveness.

##### Blocked-cyclic naming

The blocked-cyclic naming task was similar to that used in Experiment two with the following exceptions: a response deadline of 2,000 ms was implemented in order to ensure that the entire task was completed while stimulation was being administered, and participants were shown a different set of exemplars from the same categories during each session. The task lasted approximately 17 minutes.

#### Procedure

The general procedure followed that of Experiment 1: Participants were first familiarized with the images to be presented in the blocked-cyclic naming task. Immediately thereafter, they performed the verb generation task. Participants then performed the blocked-cyclic naming task, which coincided with the onset of real or sham stimulation administration. Participants completed the study after two experimental sessions, one involving real (cathodal) tDCS and one involving sham (placebo) tDCS. The order of stimulation sessions (real vs. sham) was counterbalanced across participants. The two sessions were separated by at least 7 days.

#### tDCS Stimulation Parameters

tDCS was administered using a battery-driven Magstim Eldith machine. 5 × 5 cm electrodes were placed in saline-soaked pads and secured to the scalp with a rubber headband. The montage used for the administration of tDCS (real and sham) was determined using a modeling stimulation system. The cathode was placed at F7 and the anode was placed in the middle of the forehead. This provided a maximal amount of stimulation to the LIFG with minimal stimulation to the temporal lobe; temporal stimulation could have affected the mapping from the semantic to phonological levels of word retrieval. During the real stimulation session, participants received 20 minutes of stimulation with an intensity of 1.5 mA. During this time, the entire task was completed. The sham setting on the tDCS box was used for the sham session and followed the same timing as cathodal stimulation with a 60 s ramp-up/ramp-down, which is typically thought to be sufficient to mimic the sensations of real stimulation prior to desensitization.

#### Data analysis

As in the previous experiments, trial-level naming latencies were analyzed using a linear mixed-effects model. Fixed effects included Stimulation Condition (cathodal vs. sham), Context Sequence (RU vs. UU), Half (first vs. second), and all interactions among these factors. The model included random intercepts for participants and items. Trial-exclusion criteria and all other model-fitting procedures followed those described for the preceding experiments.

Experiment 3 tested whether cathodal stimulation slowed naming of RU items both when they appeared in related blocks and after they were redistributed into unrelated blocks. These predictions were informed by the findings of Experiments 1 and 2. We therefore specified planned contrasts comparing cathodal and sham stimulation separately for RU and UU Context Sequences in the first and second half of the task. The RU contrasts tested the predicted slowing of RU items following cathodal stimulation and its persistent across halves, whereas the UU contrasts assessed stimulation effects on items presented exclusively in unrelated contexts. Because these contrasts addressed specific a prior prediction, they were evaluated regardless of the significance of the omnibus Session × Context Sequence × Half interaction.

#### Results

A total of 2.92% data points from cycles 2-4 (403) were excluded due to comprising of incorrect responses or false starts (2.65%) and equipment or microphone errors (0.27%).

Neither the Session × Context Sequence × Half interaction (*F*(1, 13262.43) = .46, *p* = .49) nor the Session × Context Sequence interaction (*F*(1, 13393.09) *=* 3.7, *p* = .055) reached statistical signficance. We nevertheless evaluated the prespecified contrasts because they tested predictions derived from Experiments 1 and 2.

In the first half of the experiment, participants named RU items more slowly under cathodal stimulation than sham stimulation, *b* = -11.24, z = −2.13, *p* < .05 Stimulation condition had no effect of naming latencies in the first half of the task for UU items, b = −11.24, z = −0.87, *p* = .39. Critically, in the second half of the task, RU items continued to elicit slower naming latencies under cathodal than sham stimulation, despite now appearing in unrelated blocks, b = −18.73, z = −3.56, *p* < .001. The corresponding contrast for UU items was not statistically significant, b = −4.96, z = −0.94, *p* = .35 (see Figure 5). Taken together, this pattern of results is consistent with Experiment 1, using a different form of NIBS and a more controlled experimental design. Complete model and contrast results are reported in Table 4.

**Figure 5.**
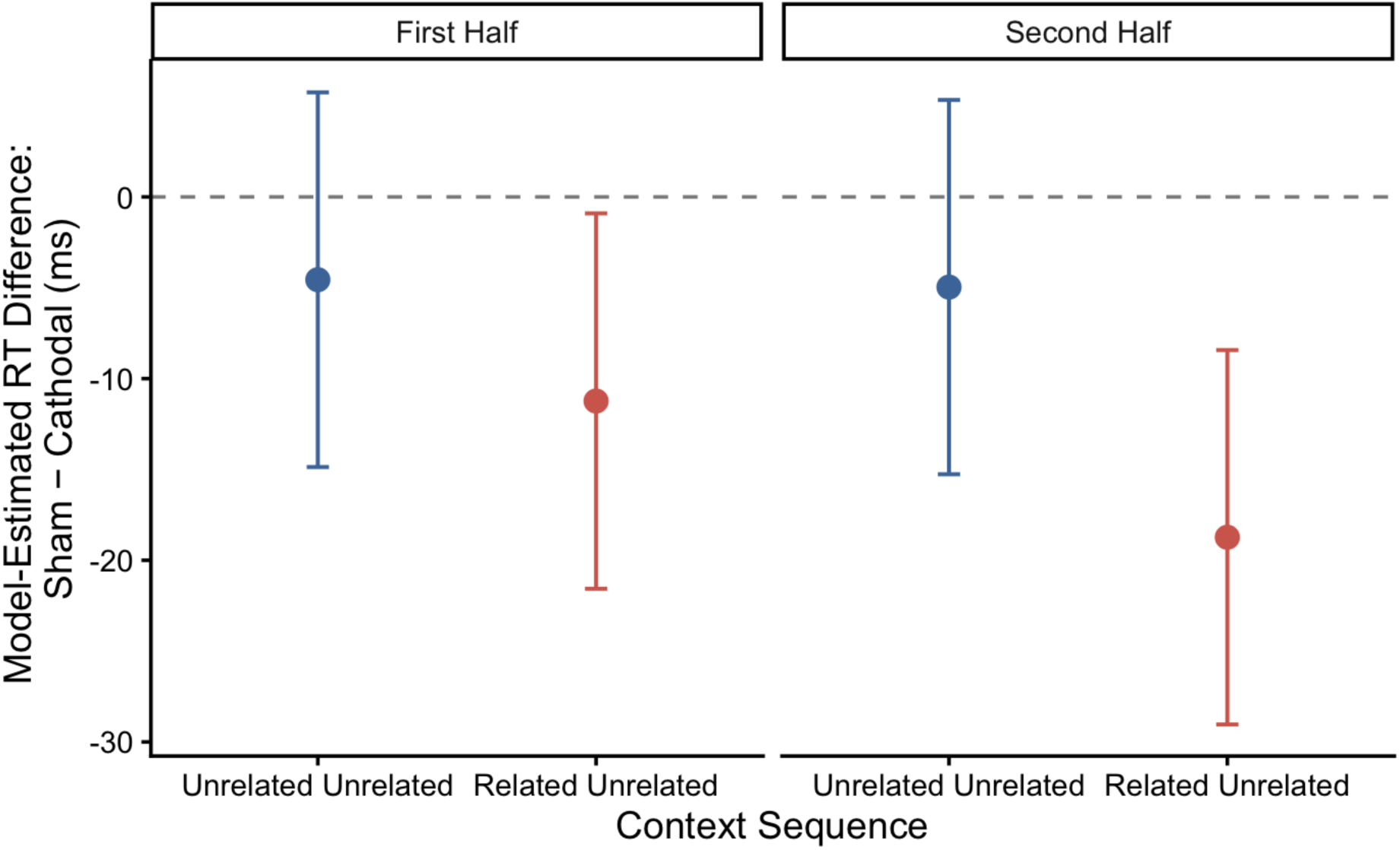
Model-estimated RT for Experiment 3. Values below the dotted line indicate faster responses during sham than cathodal.

**Table 4.** Experiment 3: Type III F Tests.

| Effect | SS | MS | $df_1$ | $df_2$ | $F$ | $p$ |
| --- | --- | --- | --- | --- | --- | --- |
| Stimulation Condition | 322508.12 | 322508.12 | 1 | 13332.00 | 14.06 | <.001 |
| Context Sequence | 676229.50 | 676229.50 | 1 | 13302.87 | 29.48 | <.001 |
| Half | 6332662.64 | 6332662.64 | 1 | 13262.43 | 276.05 | <.001 |
| Stimulation Condition ×<br>Context Sequence | 84800.92 | 84800.92 | 1 | 13393.09 | 3.70 | .055 |
| Stimulation Condition ×<br>Half | 13099.63 | 13099.63 | 1 | 13262.32 | 0.57 | .450 |
| Context Sequence ×<br>Half | 646123.02 | 646123.02 | 1 | 13262.47 | 28.17 | <.001 |
| Stimulation Condition ×<br>Context Sequence ×<br>Half | 10541.11 | 10541.11 | 1 | 13262.43 | 0.46 | .498 |

*Note.* SS = sum of squares; MS = mean square; $df_1$ = numerator degrees of freedom; $df_2$ = denominator degrees of freedom. × denotes an interaction term.

#### Experiment 3 Summary

Using a different experimental design and form of NIBS, Experiment 3 reproduced the pattern observed in Experiment 1. Relative to sham, cathodal stimulation selectively slowed items initially named in related contexts, and this effect persisted when those items were subsequently presented in unrelated contexts. In contrast, cathodal and sham stimulation did not differ for items consistently named in unrelated contexts. Thus, the effect of LIFG stimulation depended on an item’s prior related context and remained evident even after that context was removed.

## General Discussion

Across three experiments, we examined the role of the LIFG in resolving semantic interference during word selection. In Experiment 1, inhibiting the LIFG did not significantly alter semantic interference overall, but exploratory analyses revealed that its effects varied over the course of the task, potentially suggesting that the LIFG helps prevent interference from prior naming episodes disrupt future word retrieval. Experiments 2 and 3 examined this possibility more directly. In Experiment 2, semantic interference was observed when items were named in related blocks but was no longer evident after those items were redistributed into unrelated blocks, establishing a behavioral baseline in which prior semantic interference did not persist after the naming context changed. In Experiment 3, cathodal tDCS over the LIFG resulted in longer naming latencies than sham stimulation for items presented in related contexts, and this slowing remained evident even when those items were subsequently named in unrelated contexts. Together, these findings suggest that the LIFG may contribute not only to resolving immediate selection demands but also to recalibrating lexical accessibility, helping to limit the carryover of interference from prior naming episodes.

Among the proposed contributions of the LIFG to word production, competition-resolution accounts emphasize its role in selecting an appropriate response among activated alternatives, particularly when selection demands are high (Thompson-Schill et al., 1997, 1998). Evidence from aphasia supports this proposal: damage to the LIFG has been associated with greater difficulty resolving semantic interference in blocked-cyclic naming (Schnur et al., 2009; see also Harvey & Schnur, 2015). Our findings, however, suggest that disrupting this region does not uniformly increase interference across the task. In Experiment 1, the overall interference effect did not differ significantly between LIFG and vertex stimulation, although exploratory analyses revealed that stimulation effects differed across task halves—when naming context changed either from related to unrelated or vice versa. Similarly, Krieger-Redwood and Jefferies (2014) found that inhibitory (low-frequency) TMS to the LIFG did not alter semantic interference when it emerges in later naming cycles. These findings leave open how LIFG-mediated control supports selection and why its contribution may vary with the demands imposed by the task. Another account emphasizes the contribution of working memory to resolving (or minimizing) interference in blocked-cyclic naming. Belke and Stielow (2013) proposed that participants maintain the current response set and use this information to bias retrieval toward relevant items. That is, the predictable structure of the task allows participants to maintain the current response set in working memory and bias lexical activation toward task-relevant items. Accordingly, interference is constrained in part by participants’ ability to use knowledge of the response set to guide retrieval. The proposed contribution of working memory therefore depends partly on the structure of the task: repeated exposure to a predictable response set gives participants an opportunity to use that knowledge strategically to guide selection (see e.g., Belke et al., 2017). Contemporary accounts increasingly characterize the LIFG as an interface between language-specific representations and broader systems that regulate cognition according to current goals, with semantic-control functions situated adjacent to and interacting with domain-general executive systems (Jackson, 2021; Diveica et al., 2023). Maintaining and applying a task-relevant strategy may depend on working-memory processes, whereas implementing that strategy may require control over competing lexical representations. Our findings are consistent with such a hybrid account. The LIFG may support the strategic use of contextual information to guide selection and help recalibrate lexical accessibility following competition. Disruption of this process could leave previously competing items more difficult to retrieve even after the naming context changes. Consistent with this possibility, items initially named in related blocks were retrieved more slowly under cathodal than sham stimulation both before and after redistribution into unrelated blocks. Because working memory was neither manipulated nor measured directly, its contribution to this pattern remains uncertain. To test this hypothesis, future studies could introduce a concurrent working-memory load into the designs used in Experiments 2 and 3 to determine whether limiting working-memory resources increases the persistence of interference after redistribution, and whether this effect is modulated by LIFG stimulation. Working memory thus offers a possible means of supporting selection among competing words, rather than an explanation that necessarily excludes selection-related control.

Our findings are compatible with an account in which working memory and selection-related control make complementary contributions to naming. Supporting this view, Crowther and Martin (2014) found that working-memory capacity and inhibitory control were associated with different aspects of performance in blocked-cyclic naming. Working memory may help maintain contextual information that biases retrieval toward relevant responses (Belke & Stielow, 2013), while selection-related control helps resolve competition among activated words, potentially by shaping experience-dependent adjustments to semantic-to-lexical connection weights (see Thompson-Schill & Botvinick, 2006, for a related proposal). In this account, contextual information maintained in working memory could guide selection, while control exerted during selection could help recalibrate lexical accessibility for future naming. Disruption of this process could leave previously competing items more difficult to retrieve even when naming under conditions that minimize lexical-semantic competition. However, because working memory was neither manipulated nor measured directly, we cannot determine whether working-memory support for naming depends on the LIFG itself or on processes that interact with LIFG-mediated selection. Building on evidence that concurrent working-memory load increases semantic interference in blocked-cyclic naming (Belke, 2008), future studies could introduce this manipulation into the design used in Experiments 2 and 3 to determine whether limiting working-memory resources increases the persistence of interference after redistribution into unrelated contexts, and whether this effect is modulated by LIFG stimulation.

The potential interaction between task demands and LIFG-mediated control may also help explain why stimulation effects on semantic interference vary across studies. Pisoni et al. (2012) found that anodal tDCS over the LIFG reduced semantic interference in blocked-cyclic naming. Our findings following cTBS and cathodal tDCS are directionally complementary, although they do not establish a uniform relationship between stimulation polarity, cortical excitability, and behavior. By contrast, Ward et al. (2022) found no modulation of picture naming following anodal LIFG stimulation in a picture–word interference task using written or auditory distractors. This difference between paradigms may be informative. Picture–word interference manipulates competition through a distractor presented on the current trial, whereas our design explicitly examined the consequences of prior naming context on subsequent word retrieval. The contrast raises the possibility that sensitivity to LIFG disruption depends partly on the demands imposed by prior selection experience. This remains a hypothesis, however, because the studies also differed in stimulation protocols and other design features. A recent meta-analysis found no significant overall effect of TMS on semantic-control performance and highlighted methodological variability, including differences in tasks and stimulation protocols, as well as publication-bias concerns (Ambrosini et al., 2024). These findings underscore the need for direct comparisons of contextual demands while holding other design features and stimulation parameters constant.

However, within the present study, the similar patterns observed across Experiments 1 and 3 strengthen the interpretation that LIFG-mediated control influences the consequences of prior naming for subsequent retrieval. Experiment 1 used cTBS administered before naming and compared LIFG with vertex stimulation, whereas Experiment 3 used cathodal tDCS delivered during naming and compared active with sham stimulation. Despite these differences in stimulation method, timing, and control condition, both experiments yielded patterns consistent with continued difficulty retrieving items previously named in related contexts when those items appeared subsequently in unrelated contexts. Convergence across these distinct approaches makes an explanation tied solely to the timing or other features of a single stimulation protocol less likely and provides complementary support for the proposed role of the LIFG in recalibrating lexical accessibility following competition.

Although blocked-cyclic naming differs substantially from everyday language production, it may engage control processes that support retrieval in changing communicative contexts. In conversation, speakers must retrieve words in light of both what has already been said and what is relevant to the current message. Maintaining contextual information may help guide selection, while recalibrating lexical accessibility following competition could limit its consequences for subsequent retrieval. The task may therefore provide a way to examine processes whose disruption contributes to word-finding difficulties in people with aphasia (PWA). Consistent with this possibility, PWA can show relatively preserved semantic activation alongside difficulty resolving competition among closely related representations (Dyson et al., 2021). Semantic interference has been demonstrated in blocked-cyclic naming in aphasia (Schnur et al., 2006), and persistent interference has also been observed in naming with multiple distractors (van Scherpenberg et al., 2021) and without (Harvey et al., 2019; Stark et al., 2025; see also Nappo et al., 2023). These findings establish the relevance of interference resolution to aphasic naming, although whether the proposed recalibration process contributes to difficulties in everyday speech remains to be tested.

If stroke disrupts semantic control, working-memory support for selection, or their interaction, the consequences of competition during one retrieval attempt may be harder to overcome when the same or related words are encountered again, as it is thought that people with aphasia have difficulty retrieving words due to noisy lexical-semantic activation akin to that elicited by the blocked-cyclic naming task. This could contribute to retrieval failures even when the original source of interference is no longer present. Such failures need not involve production of an incorrect word: Chen et al. (2019) linked omission errors in post-stroke aphasia to impairments in semantically driven retrieval and lexical selection. Whether lingering interference contributes to these omissions or other naming errors is an open question. The potential relevance of working memory extends beyond single-word naming: Martin and Schnur (2019) associated semantic working-memory capacity with phrasal elaboration in narrative speech, and Zahn et al. (2023) demonstrated this contribution after accounting for single-word production abilities. Together, these findings motivate examining how selection-related control and working memory support retrieval across successive utterances. Future work should test whether individual differences in these abilities predict the persistence of interference in aphasia and whether interventions that strengthen adaptation to changing contexts improve subsequent naming and connected speech.

Several considerations should guide interpretation of these findings. The task-half interaction in Experiment 1 emerged from exploratory analyses, although it generated predictions that Experiments 2 and 3 were designed to examine more directly. The modest sample sizes, particularly in Experiment 3, also limit the precision of the estimated effects and the ability to detect interactions. Experiment 3 posed additional practical constraints because naming was performed during stimulation: faster response deadlines were necessary to fit the task within the stimulation window, potentially limiting the opportunity to observe prolonged retrieval under cathodal stimulation. These constraints are especially relevant when the predicted effect is a slowing of naming. Despite differences in task timing and stimulation method, the similar patterns across Experiments 1 and 3 provide promising evidence that warrants further testing in larger samples. Future studies could allow more time for responses while retaining sufficient trials within the stimulation period. The present findings also leave open the mechanism underlying the proposed recalibration of lexical accessibility and the contribution of working memory. Combining explicit manipulations of working-memory demands with physiological measures of stimulation response would help clarify both the cognitive processes involved and the effects of stimulation on LIFG function.

## Conclusion

In summary, the present study provides converging evidence that the contribution of the LIFG to word retrieval extends beyond immediate selection demands to the influence of prior naming experience on subsequent retrieval. We propose that the LIFG-mediated control helps recalibrate lexical accessibility following competition, limiting its lingering effects when words are encountered in new contexts. Working memory may support this process by maintaining contextual information that guides selection, while control during selection may shape subsequent lexical accessibility. The present findings do not establish how these contributions interact, but the similar patterns observed with cTBS and cathodal tDCS provide complementary evidence for investigating this account. For people with aphasia, difficulty recovering from prior competition could contribute to word-retrieval failures beyond the context in which that competition arose. Testing this possibility could clarify whether strengthening adaptation to changing contexts offers a useful target for language rehabilitation.

## Contributions

Laboratory for Cognition and Neural Stimulation

Study conception and design: Hamilton, Harvey, and Juhel

Acquisition of data: Harvey and Juhel

Analysis and interpretation of data: Harvey, Juhel, and Stoll

Drafting of manuscript: Harvey, Juhel, and Stoll

Critical revision: Harvey, Juhel, and Stoll

Funding: CURF Grant for Faculty Mentoring Undergraduate Research, R01DC012780 (PI Hamilton)

## Footnotes

1 Association strength and competition demand values were obtained from Snyder et al. (2011); see also Snyder & Munakata (2008).

2 No upper response-time cutoff was applied because stimulation was predicted to slow word retrieval, particularly in the semantically related condition. An upper cutoff could therefore disproportionately exclude valid responses reflecting the hypothesized stimulation effect and underestimate its magnitude.

## References

Acheson, D., & Hagoort, P. (2013). Stimulating the brain’s language network: Syntactic ambiguity resolution after TMS to the inferior frontal gyrus and middle temporal gyrus. Journal of Cognitive Neuroscience, 25(10), 1664–1677.

An, H., Bashir, S., Cha, E., Lee, J., Ohn, S., Jung, K., & Yoo, W. (2022). Continuous theta-burst stimulation over the left posterior inferior frontal gyrus induced compensatory plasticity in the language network. Frontiers in Neurology, 1–8.

Belke, E. (2008). Effects of working memory load on lexical-semantic encoding in language production. Psychonomic Bulletin & Review, 15(2), 357–363.

Belke, E. & Stielow, A. (2013). Cumulative and non-cumulative semantic interference in object naming: Evidence from blocked and continuous manipulations of semantic context. The Quarterly Journal or Experimental Psychology, 00 (0).

Costafreda, S., Fu, C., Lee, L., Everitt, B., Brammer, M., & David, A. (2006). A systematic review and quantitative appraisal of fMRI studies of verbal fluency: Role of the left inferior frontal gyrus. Human Brain Mapping, 27, 799–810.

Dell, G., Schwartz, M., Martin, N., Saffran, E., & Gagnon, D. (1997). Lexical access in aphasic and nonaphasic speakers. Psychological Review, 104(4), 801–838.

Devlin, J. T., & Watkins, K. E. (2007). Stimulating language: Insights from TMS. Brain, 130(3), 610–622.

Gervits, F., Ash, S., Coslett, H. B., Rascovsky, K., Grossman, M., & Hamilton, R. (2016). Transcranial direct current stimulation for the treatment of primary progressive aphasia: An open-label pilot study. Brain and Language, 162, 35–41.

Hallam, G., Thompson, H., Hymers, M., Millman, R., Rodd, J., Lambon Ralph, M., Smallwood, J., & Jefferies, E. (2018). Task-based and resting-state fMRI reveal compensatory network changes following damage to left inferior frontal gyrus. Cortex, 99, 150–165.

Harvey, D. & Schnur, T. (2015). Distinct loci of lexical and semantic access deficits in aphasia: Evidence from voxel-based lesion-symptom mapping and diffusion tensor imaging. Cortex, 67, 37–58.

Harvey D. & Schnur, T. (2016). Different loci of semantic interference in picture naming vs. word-picture matching tasks. Frontiers in Psychology, 7: 710.

Harvey, D., Wurzman, R., Shah-Basak, P., Faseyitan, O., Sacchetti, D., & Hamilton, R. (2016). Inhibiting the left inferior frontal gyrus via TMS has a persistent effect on naming same-category pictures. Poster (paper in prep).

Hirshorn, E., Thompson-Schill, S. (2006). Role of the left inferior frontal gyrus in covert word retrieval: Neural correlates of switching during verbal fluency. Neuropsychologia, 44(12), 2547–2557.

Howard, D., Lyndsey, N., Coltheart, M., & Cole-Virtue, J. (2006). Cumulative semantic inhibition in picture naming: experimental and computational studies. Cognition, 100, 464–482.

Ishkhanyan, B., Lange, V., Boye, K., Mogensen, J., Karabanov, A., Hartwigsen, G., & Siebner, H. (2020). Anterior and posterior left inferior frontal gyrus contribute to the implementation of grammatical determiners during language production. Frontiers in Psychology, 11, 1–13.

Kelly, H., Brady, M.C., & Enderby, P. (2010). Speech and language therapy for aphasia following stroke. Cochrane Database of Systematic Reviews, 5.

Klaus, J., & Hartwigsen, G. (2019). Dissociating semantic and phonological contributions of the left inferior frontal gyrus to language production. Human Brain Mapping, 40, 3279–3287.

Kuznetsova A, Brockhoff PB, Christensen RHB (2017). lmerTest Package: Tests in Linear Mixed Effects Models. Journal of Statistical Software 82(13), 1–26. doi:10.18637/jss.v082.i13<https://doi.org/10.18637/jss.v082.i13>.

Marangolo, P., Marinelli, C.V., Bonifazi, S., Fiori, V., Ceravolo, M.G., Provinciali, L., & Tomaiuolo, F. (2011). Electrical stimulation over the left inferior frontal gyrus (IFG) determines long-term effects in the recovery of speech apraxia in three chronic aphasics. Behavioural Brain Research, 225, 498–504.

Mattioli, F., Ambrosi, C., Mascaro, L., Scarpaza, C., Pasquali, P., Frugoni, M., Magoni, M., Biagi, L., & Gasparotti, R. (2013). Early aphasia rehabilitation is associated with functional reactivation of the left inferior frontal gyrus: A pilot study. Stroke, 45(2), 545–552.

Medaglia, J., Harvey, D., Kelkar, A., Zimmerman, J., Mass, J., Bassett, D., & Hamilton, R. (2021). Language tasks and the network control role of the left inferior frontal gyrus. eNeuro, 8(5), 1–18.

Medaglia, J., Harvey, D., White, N., Kelkar, A., Zimmerman, J., Bassett, D., & Hamilton, R. (2018). Network controllability in the inferior frontal gyrus relates to controlled language variability and susceptibility to TMS. Journal of Neuroscience, 38, 6399–6410.

Meinzer, M., Yetim, O., McMahon, K., & Zubicaray, G. (2016). Brain mechanisms of semantic interference in spoken word production: An anodal transcranial direct current stimulation (atDCS) study. Brain and Language, 157–158, 72-80.

Oppenheim, G., Dell, G., & Schwartz, M. (2010). The dark side of incremental learning: A model of cumulative semantic interference during lexical access In speech production. Cognition, 114, 227–252.

Pisoni, A., Papagno, C., & Cattaneo, Z. (2012). Neural correlates of the semantic interference effect: new evidence from transcranial direct current stimulation. Neuroscience, 233, 56–67.

Price, A., McAdams, H., Grossman, M., & Hamilton, R. (2015). A meta-analysis of transcranial direct current stimulation studies examining the reliability of effects on language measures. Brain Stimulation, 8(6), 1093–1100.

Rosen, H., Petersen, S., Linenweber, M., Snyder, A., White, D., Chapman, L., Dromerick, A., Fiez, J., & Corbetta, M. (2000). Neural correlates of recovery from aphasia after damage to left inferior frontal cortex. Neurology, 55(12), 1883–1894.

Schnur, T., Schwartz, M., Brecher, A., & Hodgson, C. (2006). Semantic interference during blocked-cyclic naming: Evidence from aphasia. Journal of Memory and Language, 54, 199–227.

Schnur, T., Schwartz, M., Kimberg, D., Hirshorn, E., Coslett, H., & Thompson-Schill, S. (2009). Localizing interference during naming: Convergent neuroimaging and neuropsychological evidence for the function of Broca’s area. PNAS, 106(1), 322–327.

Thompson-Schill, S., D’Esposito, M., Aguirre, G., & Farah, M. (1997). Role of left inferior prefrontal cortex in retrieval of semantic knowledge: A reevaluation. Proc. Natl. Acad. Sci., 94, 14792–14797.

Thompson-Schill, S., Swick, D., Farah, M., D’Esposito, M., Kan, I., & Knight, R. (1998). Verb generation in patients with focal frontal lesions: A neuropsychological test of neuroimaging findings. Proc. Natl. Acad. Sci., 95, 15855–15860.

Thompson-Schill, S. & Botvinick, M. (2006). Resolving conflict: A response to Martin and Cheng (2006). Psychonomic Bulletin & Review, 13, 402–408.

Tyler, L., Marslen-Wilson, W., Randall, B., Wright, P., Devereux, B., Zhuang, J., Papoutsi, M., Stamatakis, E. (2011). Left inferior frontal cortex and syntax: function, structure, and behaviour in patients wtih left hemisphere damage. Brain, 134(2), 415–431.

van Oers, C., Vink, M., van Zandvoort, M., van der Worp, H., de Haan, E., Kappelle, L., Ramsey, N., & Dijkhuizen, R. (2010). Contribution of the left and right inferior frontal gyrus in recovery from aphasia. A functional MRI study in stroke patients with preserved hemodynamic responsiveness. NeuroImage, 49(1), 885–893.

Winhuisen, L., Thiel, A., Schumacher, B., Kessler, J., Rudolf, J., Haupt, W., & Heiss, W. (2005). Role of the contralateral inferior frontal gyrus in recovery of language function in poststroke aphasia: A combined repetitive transcranial magnetic stimulation and positron emission tomography study. Stroke, 36(8), 1759–1763.

Winhuisen, L., Thiel, A., Schumacher, B., Kessler, J., Rudolf, J., Haupt, W., & Heiss, W. (2007). The right inferior frontal gyrus and poststroke aphasia: A follow-up investigation. Stroke, 38(4), 1286–1292.

Xie, X., & Myers, E. (2018). Left inferior frontal gyrus sensitivity to phonetic competition in receptive language processing: A comparison of clear and conversational speech. Journal of Cognitive Neuroscience, 30(3), 267–280.

Zhu, Z., Zhang, J., Wang, S., Xiao, Z., Huang, J., & Chen, H. (2009). Involvement of left inferior frontal gyrus in sentence-level semantic integration. NeuroImage, 47(12), 756–763.

