## Supplementals for "The Role of the Left Inferior Frontal Gyrus in Resolving Immediate and Carryover Semantic Interference During Picture Naming"

**Table S1**

*Stimulus Items for Experiment 1*

| Category | Mixed (unrelated) set | | | | | |
|---|---|---|---|---|---|---|
| | 1 | 2 | 3 | 4 | 5 | 6 |
| *List 1a* | | | | | | |
| Animals | cat | cow | dog | horse | pig | rabbit |
| Appliances | blender | mixer | washing machine | stove | TV | clock |
| Food | donut | bacon | cupcake | cookie | bread | hot dog |
| Furniture | chair | crib | desk | wardrobe | mattress | pillow |
| Instruments | banjo | violin | harp | piano | saxophone | flute |
| Tools | nail | shovel | saw | screwdriver | hammer | wrench |
| *List 1b* | | | | | | |
| Clothing | sandal | pants | jacket | shirt | hat | dress |
| Fruit | lime | blueberry | watermelon | raspberry | cherries | pineapple |
| Office supplies | envelope | highlighter | notebook | pencil | scissors | stapler |
| Sea creatures | dolphin | octopus | seal | shark | crab | whale |
| Utensils | chopsticks | fork | knife | ladle | measuring cup | spoon |
| Vehicles | plane | bus | golf cart | bicycle | tractor | jeep |
| *List 2a* | | | | | | |
| Animals | elephant | giraffe | lion | monkey | tiger | zebra |
| Appliances | cellphone | fan | toaster | radio | iron | vacuum |
| Food | rice | bagel | pizza | spaghetti | ham | popcorn |
| Furniture | bed | rug | stool | lamp | dresser | table |
| Instruments | accordion | xylophone | drum | guitar | trumpet | clarinet |
| Tools | rake | ladder | bucket | paintbrush | axe | screw |
| *List 2b* | | | | | | |
| Clothing | robe | T-shirt | sweater | sock | shoe | boot |
| Fruit | plum | pear | lemon | tomato | apple | strawberry |
| Office supplies | calculator | computer | paperclip | staples | printer | pen |
| Sea creatures | eel | starfish | turtle | lobster | seahorse | snail |
| Utensils | bowl | cup | spatula | plate | rolling pin | thermos |
| Vehicles | car | boat | motorcycle | train | helicopter | van |

*Note.* Each row lists the six items forming a related (same-category) set; each numbered column forms a mixed (unrelated) set containing one item from each category in that sublist. Each participant saw both lists, one per session, with list order counterbalanced across participants. Each list was divided into two sublists (e.g., List 1a and List 1b). Within a session, items in one sublist appeared in related sets during the first half and unrelated sets during the second half. Items in the other sublist followed the opposite pattern, appearing in unrelated sets first and related sets second.

**Table S2**

**Table S2**

*Stimulus Items for Experiments 2 and 3*

| | Mixed (unrelated) set | | | | | |
|---|---|---|---|---|---|---|
| Category | 1 | 2 | 3 | 4 | 5 | 6 |
| *Group 1* | | | | | | |
| Animals | elephant | giraffe | lion | monkey | tiger | zebra |
| Appliances | cellphone | fan | toaster | radio | iron | vacuum |
| Food | rice | bagel | pizza | spaghetti | ham | popcorn |
| Furniture | bed | rug | stool | lamp | dresser | table |
| Instruments | accordion | xylophone | drum | guitar | trumpet | clarinet |
| Tools | rake | ladder | bucket | paintbrush | axe | screw |
| *Group 2* | | | | | | |
| Clothing | robe | T-shirt | sweater | sock | shoe | boot |
| Fruit | plum | pear | lemon | tomato | apple | strawberry |
| Office supplies | calculator | computer | paperclip | staples | printer | pen |
| Sea creatures | eel | starfish | turtle | lobster | seahorse | snail |
| Utensils | bowl | cup | spatula | plate | rolling pin | thermos |
| Vehicles | car | boat | motorcycle | train | helicopter | van |

*Note.* Each row lists the six items forming a related (same-category) set; each numbered column forms a mixed (unrelated) set containing one item from each category in that group. Items were divided into two groups. For each participant, one group was presented in related sets and the other in unrelated sets, with group assignment counterbalanced across participants. In the second half of the experiment, the related sets were rearranged into unrelated sets, while the unrelated sets remained unrelated.
